# KATANIN is required for the formation of complete preprophase bands and subsequent division positioning of subsidiary mother cells in maize

**DOI:** 10.64898/2026.05.17.725781

**Authors:** Stephanie E. Martinez, Carolyn G. Rasmussen

## Abstract

Microtubule dynamics underpin cell division, movement, and growth in eukaryotic organisms. KATANIN p60 is a microtubule-severing protein that promotes proper cell elongation and division. In plants, division positioning is initiated in late G2 via the formation of a microtubule structure called the preprophase band (PPB). Maize *p60* mutants have defects in microtubule severing and form abnormal PPBs in symmetrically dividing cells, including both incompletely assembled and misoriented PPBs. Here, we show that an asymmetric division required to generate stomatal complexes in maize *p60* mutants have normally positioned but often incompletely formed PPBs. Incompletely formed PPBs lead to misoriented divisions and nuclear positioning defects in *p60* mutants.

## Introduction

Microtubule dynamics and organization are important for many cellular processes, including division, elongation, and differentiation (Hashimoto, 2015; Horio and Murata, 2014). Microtubule severing is essential for microtubule reorganization and is performed in many eukaryotes by AAA-type (ATPases Associated with diverse cellular Activities) ATPases like KATANIN, a complex composed of p60 and p80 subunits. The p80 subunit recruits the p60 to microtubules for p60-mediated ATP-dependent microtubule severing (Deinum et al., 2017; Hartman and Vale, 1999; Hartman et al., 1998; McNally and Vale, 1993; Wang et al., 2017; Zhang et al., 2013). In animals, p60 microtubule severing is important for completion of cytokinesis (Matsuo et al., 2013), meiosis (Dunleavy et al., 2023; Lynn et al., 2021; Srayko et al., 2000) and many other processes (Ghosh et al., 2012). In plants, *p60* mutants have aberrant microtubule organization, leading to shorter, wider cells and small organs (Attrill and Dolan, 2024; Bichet et al., 2008; Burk et al., 2001; Komis et al., 2017; Komorisono et al., 2005; Luptovčiak et al., 2017; Martinez et al., 2026; Panteris and Adamakis, 2012; Wang et al., 2021). In *Zea mays* (maize), p60 is encoded by two genes, *Discordia3a (Dcd3a)* and *Dcd3b*, both of which play mostly redundant roles in growth and fertility (Martinez et al., 2026).

In grasses, such as maize and *Brachypodium distachyon*, stomatal complexes have two guard cells flanked by two subsidiary cells (Berg and Raissig, 2025; Gray et al., 2020; Stebbins and Shah, 1960). Subsidiary cells are formed from an asymmetric division that generates a small subsidiary cell directly in contact with the guard cell mother cell and a pavement cell. The division positioning is very consistent in these cells (Guo et al., 2021; McKown and Bergmann, 2020). Before the onset of cell division, the nucleus polarizes adjacent to the guard mother cell for the asymmetric division of the subsidiary mother cell (Facette et al., 2019). Nuclear polarization is mediated by the proper localization of BRICK1, which recruits Leucine-rich-repeat receptor-like kinase PANGLOSS2 (PAN2; reviewed in (Zhou et al., 2026). PAN2 recruits the WEAK CHLOROPLAST MOVEMENT UNDER BLUE LIGHT 1 (WEB1)/PLASTID MOVEMENT IMPAIRED 2 (PMI2)-RELATED (WPR) proteins, which interact directly with F-actin (Nan et al., 2023). PAN2 also recruits PANGLOSS1 (PAN1) and plant-specific Rho-of-plants (ROP) GTPases to enhance actin patch accumulation at the interface between the subsidiary mother cell and the guard cell (Facette et al., 2015; Humphries et al., 2011; Zhang et al., 2012). The maize LINC KASH SINE-LIKE2 (MLKS2) mediates the polarized movement and tethering of the nucleus to the actin patch (Ashraf et al., 2023). Then, during late G2, the PPB, a cortical ring of microtubules and actin filaments, forms and marks the division site, or the location of the division after the completion of cytokinesis (Figure 1A).

**Figure 1.**
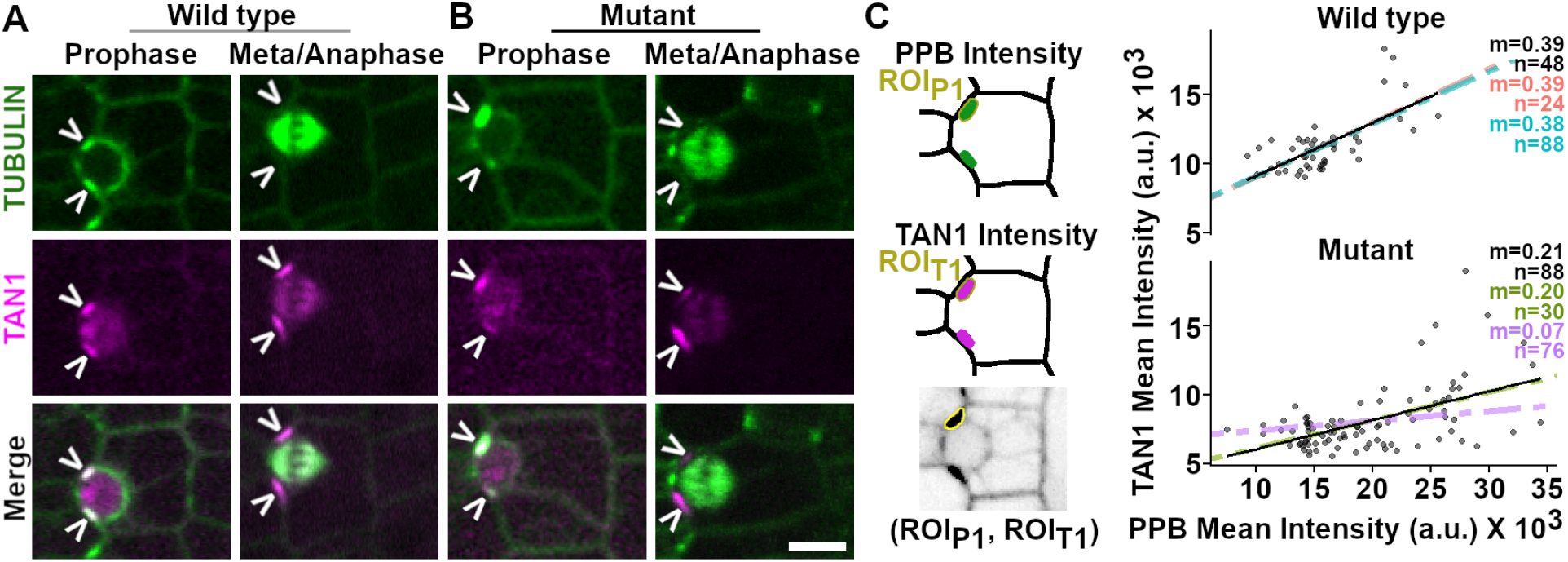
TAN1 is a proxy for PPB accumulation. A, B) Representative micrographs of asymmetrically-dividing subsidiary cells expressing CFP-TUBULIN (microtubules) and TAN1-YFP (magenta) for A) wild-type cells (*dcd3a-2/+; dcd3b-1/+*) with equivalent accumulation of both microtubules and TAN1-YFP and B) *p60* mutant cells (*dcd3a-2; dcd3b-1)* with uneven accumulation of both TAN1-YFP and microtubules at each side of the division site. White arrowheads indicate the division site. All images are the same size; scale bar = 10 μm. C) Schematic showing the division site where fluorescent intensities were measured as well as a graph comparing relative TAN1 and PPB mean intensity from cells in late G2/prophase. Three plants per genotype were analyzed. Raw data from one plant consists of black dots corresponding to individual fluorescence intensity measurements (with a black line of best fit); colored dotted lines = best fit of 2 other plants. m = slope, n = number of division site fluorescence intensities measured.

The PPB recruits a suite of proteins, of which a small subset remains at the division site after PPB disassembly, including the microtubule-binding protein TANGLED1 (Bellinger et al., 2023; Livanos and Müller, 2019; Martinez et al., 2017; Rasmussen et al., 2011; Smith et al., 2001; Uyehara and Rasmussen, 2023; Uyehara et al., 2024; Walker et al., 2007). Live-cell timelapse imaging demonstrated that TAN1 localization at the division site during later stages of mitosis indicated prior PPB accumulation. Therefore, TAN1 division site localization is an excellent proxy for earlier PPB accumulation (Uyehara et al., 2024).

In maize, *Marchantia polymorpha*, and Arabidopsis, *p60* mutants have a combination of incompletely formed PPBs, misoriented PPBs and division plane orientation defects (Attrill and Dolan, 2024; Komis et al., 2017; Martinez et al., 2026; Ovečka et al., 2020; Panteris et al., 2011; Webb et al., 2002). Following PPB disassembly, the mitotic spindle forms in metaphase and separates chromosomes in anaphase (Figure 1A). In telophase, the phragmoplast, an anti-parallel microtubule array, expands towards the division site and facilitates the formation of the cell plate, which will mature into the cell wall that separates the two daughter cells (Müller and Jürgens, 2016; Rasmussen and Bellinger, 2018; Samuels et al., 1995). Phragmoplasts often grow with arrow-head shaped aberrantly oblique leading edges in p60 mutants (Komis et al., 2017; Martinez et al., 2026; Panteris et al., 2011). Near the cell cortex, cortical telophase microtubules, which are stabilized by TAN1 or other division-site localized proteins, are added into the phragmoplast to direct phragmoplast positioning (Bellinger et al., 2023). In *p60* mutants, delayed microtubule re-organization leads to defects in cell wall formation during cytokinesis (Panteris et al., 2021).

In symmetric divisions, maize *p60* mutants (*dcd3a-2; dcd3b-1* double mutants) formed both incomplete and misoriented PPBs. Incomplete PPBs, containing uneven or one-sided microtubule accumulations at the expected division site, did not lead to misoriented divisions. Only rarely occurring (5%) misoriented PPBs, containing microtubule accumulations not at the division site but in another aberrant location, led to misoriented symmetric divisions in *p60* mutants. Subsidiary cells were occasionally misoriented ∼15% (Martinez et al., 2026). We hypothesized that, similar to symmetric divisions, misoriented asymmetric divisions would be caused by misoriented PPBs. In contrast to our hypothesis, there were no misoriented PPBs in asymmetric subsidiary cell divisions. Instead, incomplete PPBs generated misoriented divisions and mispositioned nuclei.

## Results and Discussion

To determine when defects arise during asymmetric division, we used live-cell imaging of *p60* mutants (*dcd3a-2; dcd3b-1* homozygous double mutants) and wild-type siblings (*dcd3a-2/+; dcd3b-1/+*) expressing CFP-TUBULIN and TAN1-YFP. First, we assessed PPB positioning in subsidiary cell divisions. There were neither misoriented PPBs nor cells completely lacking PPBs in either mutants or wild-type plants (n = 3 plants per genotype, ≥ 98 cells analyzed).

Next, we assessed whether PPBs were normal or incomplete. We defined normal PPBs as those with uniform fluorescence intensities on both sides of the division site at the midplane of the cell with < 30% difference (Figure 1A). Normal PPBs were observed 97% of the time in wild-type plants (n = 95/98 cells from 3 plants) and 73% of the time in *p60* mutants (n = 102/139 cells from 3 plants). The rest of the PPBs in the *p60* mutant were either uneven (∼23%, n = 32/139 cells with 30-75% fluorescence intensity differences at the division site, Figure 1B) or one-sided (∼4%, n =5/139 cells with fluorescence intensity differences > 75%). These data indicated that PPBs in these asymmetric divisions were not misoriented, but instead incompletely formed.

To confirm that TAN is a reliable proxy for prior PPB accumulation at the division site, we directly compared TAN1 localization with the microtubule PPB. TAN1 localization mirrored PPB type ∼97% of the time in wild-type siblings and ∼96% of the time in double mutants (n = 81/84 wild-type cells; n = 44/46 cells mutant cells; 3 plants each genotype). Similar to previous research (Uyehara et al., 2024), there was a positive correlation between TAN1 and PPB mean fluorescence intensity in late G2/prophase (Figure 1C; R^2^ ≥ 0.3 for 3 plants each genotype). Further, uneven TAN1 fluorescence intensities in *p60* mutant cells during telophase occurred with similar frequency to uneven PPB accumulations during G2/prophase (37%, n = 11/29 cells from 3 mutant plants vs. 23% uneven PPBs, p =0.09 Chi-square test). Together, these data indicate that TAN1 is an accurate marker of the division site previously marked by the PPB.

Next, we assessed TAN1 accumulation in telophase cells to determine whether incomplete PPBs caused misoriented asymmetric divisions. Misoriented phragmoplast ends were strongly correlated with local reduced TAN1 accumulation at the division site in the *p60* mutant. In wild-type siblings, phragmoplasts were correctly oriented and had a TAN1 fluorescence intensity ratio at the division site close to 1, indicating a prior normal PPB (Figure 2A, C). In *p60* mutants, we observed variable TAN1 fluorescence ratios during telophase indicative of normal, uneven, and one-sided PPBs. When TAN1 accumulation was only on one side of the division site, the phragmoplast was oriented on that side, and misoriented on the other side (Figure 2B, C, TAN1 fluorescence ratio ≤ 0.4, n = 6/6 one-sided TAN1 cells) In addition, when there was uneven TAN1 accumulation, correct phragmoplast orientation was correlated with higher TAN1 accumulation (n = 9/9). Sometimes, uneven TAN1 accumulation did not lead to a misoriented phragmoplast (n = 4/9 cells) (Figure 2C). This shows that one-sided PPBs, and occasionally uneven PPBs, cause misoriented asymmetric divisions in the *p60* mutant. Similarly, another mutant which forms incomplete (one-sided and uneven) PPBs, called *discordia1* (*dcd1*), had misoriented asymmetric divisions that occurred due to local absence of the PPB and lack of TAN1 accumulation (Uyehara et al., 2024). Taken together, this indicates that incomplete PPBs lead to asymmetric division positioning defects. More incomplete PPBs are observed during symmetric divisions than asymmetric divisions in the *p60* mutant (41% vs 27%, respectively). However, incomplete PPBs do not cause misoriented symmetric divisions (Martinez et al. 2026). This indicates a different, and perhaps more critical, role for the PPB in asymmetric division plane positioning.

**Figure 2.**
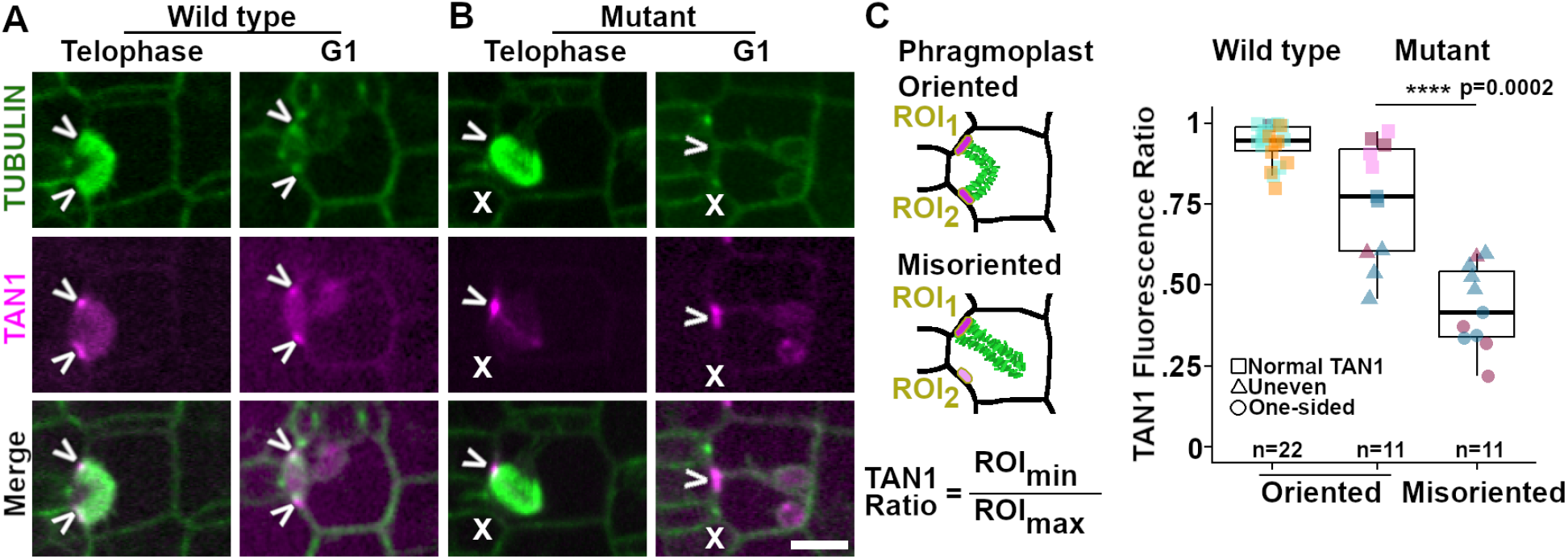
Incomplete PPB accumulation leads to subsidiary cell asymmetric defects. A, B) Representative micrographs of asymmetrically-dividing subsidiary mother cells in telophase/cytokinesis and early G1 expressing CFP-TUBULIN (microtubules) and TAN1-YFP (magenta). White arrowheads indicate the division site, while the white Xs show division sites lacking TAN1. A) Representative micrograph of wild-type cells with even TAN1 fluorescence at both sides of the division site and oriented phragmoplasts and B) *p60* mutants with one-sided TAN1 accumulation and a misoriented phragmoplast. All images are the same size; scale bar = 10 μm. C) TAN1 fluorescence ratio for cells with oriented phragmoplasts: Wild type (0.94 ± 0.01), Mutant (0.76 ± 0.06); and misoriented phragmoplasts: Mutant (0.43 ± 0.04). No misoriented phragmoplasts were observed in wild type. Three plants from each genotype were analyzed; each color = individual plant. Wilcoxon rank sum test p-values and cell counts shown.

Aberrant PPBs in symmetric divisions cause nuclear positioning defects in *p60* mutants. To determine if nuclear positioning is influenced by the PPB in asymmetric divisions, the distance of the nucleus relative to the maximum PPB accumulation was measured (Figure 3). Similar to wild-type siblings, *p60* mutant cells with normal PPBs had a mean nuclear position of ∼0.5, indicating that the nucleus was evenly-centered between the two PPB accumulations (Figure 3A, C). However, in double mutant cells with incomplete PPBs, the nucleus was closer to the maximum PPB accumulation (Figure 3B, C, mean nucleus position ∼0.4). This indicates that nuclei in mutants were closer to the maximum PPB accumulation.

**Figure 3.**
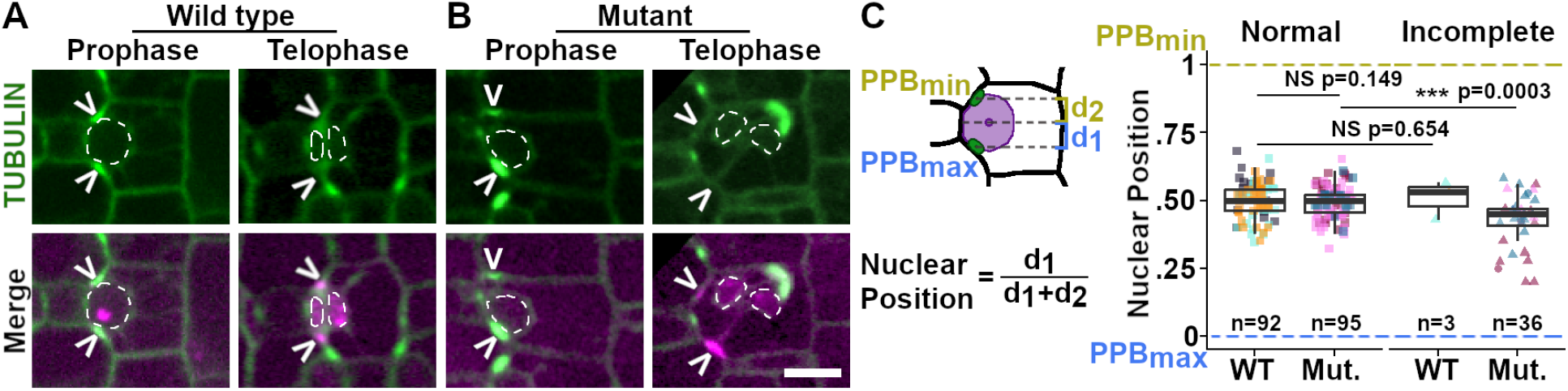
Incomplete PPBs influence nuclear positioning towards microtubule accumulation. A, B) Representative micrographs of cells expressing CFP-TUBULIN (green) and TAN1-YFP (magenta) during prophase and telophase in A) wild-type siblings and B) *p60* mutants. White arrowheads indicate the division site, nuclei are outlined with dotted lines. All images are the same scale; scale bar = 10 μm. C) Normalized nuclear position in cells with normal PPBs (Wild type: 0.50 ± 0.01; Mutant: 0.49 ± 0.01) and incomplete PPBs (Wild type: 0.51 ± 0.04; Mutant: 0.43 ± 0.02). Three plants from each genotype were analyzed; each color = individual plant. For shapes: square = normal PPB, triangle = uneven PPB, circle = one-sided PPB. Wilcoxon rank sum test p-values and cell counts shown.

Nuclear polarization, or the movement of the nucleus towards the cell wall adjacent to the guard mother cell, occurs before PPB positioning in asymmetric divisions (Hazelwood et al., 2025).Nuclear polarization defects cause misoriented PPBs that later generate misoriented asymmetric divisions, as seen in the maize linc kash sine-like2 (*mlks2*) mutant (Ashraf et al., 2023; Gumber et al., 2019). An additional maize mutant with subsidiary mother cell asymmetric division defects, *aladin1*, encodes a component of the nuclear pore complex. Whether the defect is due to PPB or nuclear positioning, or something else, is currently unknown (Best et al., 2021). Even though misoriented divisions were observed in *p60* mutants, it was not due to a nuclear polarization defect because nuclei polarized correctly and PPBs were positioned at the expected division sites (Figures 2, 3). This indicates that misoriented divisions in the *p60* mutant are independent of nuclear polarization prior to PPB formation. It is currently unclear whether nuclear positioning defects after PPB formation contribute to division plane positioning but there is no evidence currently suggesting that it does.

## Materials and Methods

### Plant Growth Conditions

Plants were grown under standard greenhouse conditions (32°C setpoint) with 4 hours of supplemental lighting (∼400 µEm-2s^-1^) in 1 L pots with soil composed of 50% bark, 20% peat, 20% medium vermiculite, and 10% perlite with calcium nitrate (75 ppm N and 90 ppm Ca), magnesium nitrate (50 ppm N and 45 ppm Mg), and Osmocote Classic 3-4M (NPK 14-14-14 %, AICL SKU#E90550).

### Genotyping

DNA was extracted from leaves collected at 2 weeks-old using a standard protocol (Lunde, 2018; Mills et al., 2020). Mutant alleles (*dcd3a-2, dcd3b*) and transgenes (CFP-β-TUBULIN, TAN1-YFP, (Mohanty et al., 2009)) were genotyped using the primers and/or restriction enzymes specified in (Martinez et al., 2026).

### Live-cell imaging of the asymmetric subsidiary mother cell divisions

Plants expressing CFP-TUBULIN and TAN1-YFP were grown under standard greenhouse conditions for 4 weeks before dissecting to the youngest emerging leaf (ligule length < 2 mm). Samples were cut 0.4-0.5 cm from the ligule and mounted with water into a rose chamber (Rasmussen, 2016). Images were obtained of the abaxial side of the leaf with a Hamamatsu 9100c EM-CCD camera using a Yokogawa W1 spinning disk microscope with Nikon Eclipse TE inverted stand. Solid-state Obis lasers (40-100 mW) of 445 nm or 514 nm with 480/40 or 540/30 nm emission filters (Chroma Technology) were used for CFP-TUBULIN and TAN1-YFP, respectively. Using a water immersion objective (60X/1.2 NA), an ASI Piezo Z stage was used to collect Z-stacks of 1-2 μm step size in Micromanager-1.4 using a 3-axis DC controller. During image acquisition, unbiased imaging was performed by tiling across the leaf sample.

### PPB and TAN1 Fluorescence Intensity Measurements

Using images taken as described above, the mean fluorescence intensity of the PPB (CFP-TUBULIN) and TAN1-YFP was obtained by outlining the PPB at the midplane of the cell using the polygon tool in FIJI. Mean gray value of the same ROI was measured in the corresponding CFP-TUBULIN and TAN1-YFP channels (Figure 1C). For cells in telophase, TAN1-YFP at the division site was outlined at the midplane of the cell (Figure 2C). For one-sided PPB or TAN1-YFP accumulations, a polygon was outlined at the other expected division site.PPB and TAN1 accumulations in a cell were classified using the following fluorescence intensity percent differences: normal < 30%, uneven 30-75%, one-sided > 75%.

### Nuclear Positioning Analysis

Using images of cells in late G2/prophase expressing CFP-TUBULIN, the nuclear and PPB centroids at the mid-plane of a cell were obtained in FIJI. A straight line was drawn through the centroids and the distance between the lines was measured (Figure 3C). For cells with one-sided PPBs, ROIs were traced at the expected location of the PPB. The normalized nuclear position was obtained by dividing the distance of the nucleus to the PPB with maximum fluorescence by the total distance between the PPBs (Figure 3C).

### Figure Preparation, Graphs, and Statistics

Gnu Image Manipulation Program (GIMP, v3.0.4, https://www.gimp.org/) was used for figure preparation. If applicable, levels were adjusted linearly and images were enlarged or rotated with no interpolation. Graphing and statistics were done in RStudio using the tidyr (Wickham et al., 2024), ggplot2 (Wickham, 2016), paletteer (Hvitfeldt, 2021), and ggprism (Dawson, 2024) packages.

## Acknowledgements

We thank Audrey Choi for some image acquisition during Summer 2022. We also thank the other members of the Rasmussen lab for helpful feedback. This work was supported by NSF grants.

## Author Contributions

Conceived and designed the analysis: CGR and SEM

Collected the data: SEM

Performed the analysis: SEM

Wrote the paper: SEM and CGR

Revised and reviewed the paper: SEM and CGR

Funding acquisition: SEM and CGR

